# Hospital use of antibiotics as the main driver of infections with antibiotic-resistant bacteria – a reanalysis of recent data from the European Union

**DOI:** 10.1101/553537

**Authors:** Matthias M. Fischer, Matthias Bild

## Abstract

Antimicrobial resistance in bacteria causes significant morbidity worldwide. The development and acquisition of resistance to antibiotics is believed to primarily develop under the selective pressure of widespread antibiotic use in humans, however antimicrobial usage in livestock has been proposed as additional, if not principal, driver of antibiotic resistance. In this work, we correlate recent data from the European Union on antibiotic resistance rates with data on antibiotic usage in the primary care and hospital sector and data on veterinary antimicrobial consumption across the individual member states. We quantify the strength of these different potential drivers of antimicrobial resistance in order to compare their biological importance. We found that the correlation between antibiotic use in the hospital sector and antibiotic resistance rates is significantly higher than the correlation between resistance rates and any of the other two predictors. This suggests increased antibiotic use in hospitals as the main driver of the development of antibiotic resistances and necessitates further research on and a re-evaluation of the risks associated with antibiotic use in human and veterinary medicine.

## 1 Introduction

Microbes are able to develop or acquire resistance towards antibiotic substances they are exposed to, which becomes problematic in case of bacteria that cause severe infectious diseases in humans (Holmes et al., 2016). This has become particularly pervasive due to a general and costly failure to discover fundamentally new classes of antibiotic substances and a change of focus of industrial research towards drugs aiming to prevent or mitigate non-infectious diseases instead. Consequently, in 2014 the World Health Organization declared that the problem of increasing antibiotic resistance rates *threatens the achievements of modern medicine* and that a *post-antibiotic era – in which common infections and minor injuries can kill – is a very real possibility for the 21st century* (Nathan and Cars, 2014). Therefore, understanding the main drivers of the development of microbial resistances towards antibiotic substances is important from a perspective of public health.

There is a general consensus that widespread use of antibiotics in humans leads to higher rates of resistance by placing a selective pressure on bacterial populations which favours the emergence of drug-resistant strains (Shallcross and Davies, 2014). For example, this correlation has been demonstrated in a recent study by *Cusini* and colleagues for antibiotic resistance rates of *Escherichia coli* and *Klebsiella pneumoniae* across eighteen departments of a hospital in Berne, Switzerland (Cusini et al., 2018). Although the use of antibiotics in hospitals only accounts for a small fraction of the overall antibiotic use, it significantly influences resistance rates because hospitals can be considered concentrated and fertile breeding grounds for newly emerging drug-resistant bacterial strains (Shallcross and Davies, 2014). A similar study from 2005 demonstrated a correlation between outpatient antibiotic use across twenty-six countries of the European Union during the years of 1997 to 2002 and observed antibiotic resistance rates (Goossens et al., 2005).

The same mechanism of selective pressure leading to higher resistance rates exist in agricultural facilities as well as repeatedly pointed out in the literature (Witte, 1998; Teuber, 2001; Heuer et al., 2011). Direct transmission of such resistant strains from agricultural facilities onto humans has been repeatedly observed, for a number of examples refer to Shea (2003). However, the direct correlation between antibiotic usage rates in agriculture and population level incidence of infections with drug resistant bacterial strains does not appear to have been studied yet.

A quantitative comparison of the effect sizes which the proposed mechanisms exert at the population-level infection rates with antibiotic-resistant bacterial strains is still missing in the literature. Hence, in this work we re-analyse recent datasets from the European Union on antibiotic resistance rates and on the use of antibiotic substances in the primary care sector, in the hospital sector and in the field of veterinary medicine in order to assess and compare the relative importance of these three potential drivers.

## 2 Materials and methods

### 2.1 Data sources

Data on antibiotic resistance rates were extracted from the 2017 annual report of the European Antimicrobial Resistance Surveillance Network (EARS-Net) by the European Centre for Disease Prevention and Control (European Centre for Disease Prevention and Control, 2018b). Data on antibiotic usage in the primary care sector and the hospital sector have been taken from the 2017 annual epidemiological report by the European Centre for Disease Prevention and Control (ECDC) (European Centre for Disease Prevention and Control, 2018a). Data on yearly veterinary medical antibiotic usage have been extracted from the 2016 European surveillance of veterinary antimicrobial consumption report by the European Medicines Agency (EMA) which monitors the overall sales of antibiotic substances per country (European Medicines Agency, 2018).

### 2.2 Statistical analysis

We only examined the datasets for *E. coli* and *K. pneumoniae*, because these provided data for the highest number of different antimicrobial classes (four or three classes of antibiotics, respectively). In contrast, the datasets for *P. aeruginosa*, *Acinetobacter* sp. and *S. pneumoniae* only reported data for two, two and one class of antimicrobial agents respectively, so they were excluded from further analyses due to reasons of statistical power. Because the rates of resistance of the different bacterial species were reported for different classes of antimicrobial agents, we analysed the data for the two bacterial species individually.

All statistical analyses were carried out in R version 3.3.1 (R Core Team, 2017). Both fitted model were generalised mixed-effects linear models with binomially distributed errors fitted with the `glme` command from the R package `lme4` (Bates et al., 2015), describing the rates of bacterial antibiotic resistance as response variable. We included yearly antibiotic usage in the primary care sector, yearly antibiotic usage in the hospital sector and yearly veterinary medical antibiotic usage as continuous fixed-effect predictor variables. Because the two EU reports on antibiotic consumption contained data with differing units, we converted the values of antibiotic use in humans from daily defined doses (DDD) per thousand inhabitants and day to antibiotic consumption in milligram per kilogram biomass and year, the same unit used in the annual report on veterinary medical antibiotic usage. To do so, we assumed that one DDD equals an average dosis of 1000 mg, following WHO guidelines for surveillance of antimicrobial use, and a mean human body mass of 60 kg (Nair and Jacob, 2016). The respective antibiotic class was included as random-intercept effect in order to account for individual differences in resistance rates between these classes. Previous research has demonstrated significant effects of the population density and the mean local temperature (Bruinsma et al., 2003; MacFadden et al., 2018) on rates of antimicrobial resistance. However, because it is not clear if these factors contribute linearly and additively to the likelihood of antimicrobial resistance, we decided to not include them as covariates of our models. Instead, we included the factor ‘country’ as additional random-intercept effect in order to account for such individual differences between the examined countries. This way, we were also able to account for other, yet undiscovered differences between the individual countries.

The assumptions of the fitted models were checked by visual inspection of diagnostic plots. In case of the model for *K. pneumoniae*, one entry (Fluoroquinolone usage and resistance rates in Lithuania) had to be removed as it was a severe outlier violating the normality assumption of the model. After fitting the models, we compared the actual resistance rates with the rates predicted by the fitted models in order to judge their goodness of fit. In order to check for significance of the different predictor variables, we performed chi-squared likelihood ratio tests on type II sum of squares using the `Anova` command from the R package `car` (Fox and Weisberg, 2011). Partial correlation coefficients of the three fixed-effect predictors were compared in order to quantify their relative importance. In order to visualise the effects of the three predictor variables, we generated model predictions and 0.65 confidence intervals of antibiotic resistance rates for varying levels of antibiotic use in representative countries and for representative antibiotic classes using the `predictInterval` function from the `merTools` R package (Knowles and Frederick, 2018).

## 3 Results

### 3.1 Model assumptions, goodness of fit

Both fitted models did not show any violation of the implicit model assumption of normally distributed residuals (*E. coli*: Shapiro-Wilk-Test, *W* = 0.98307, *p* > 0.05; *K. pneumonia*: Shapiro-Wilk-Test, *W* = 0.9788, *p* > 0.05). Diagnostic plots of the residuals can be found in Appendix A, as well as plots of the residuals as a function of fitted model values, showing a constant variance of the residuals regardless of the fitted values. Both models showed a high goodness of fit with *R*^2^ values of 0.983 (*E. coli*) and 0.990 (*K. pneumonia*). Diagnostic plots comparing fitted with actual values can be found in Appendix B.

### 3.2 Comparison of significance and partial correlation coefficients

#### 3.2.1 E. coli

The final dataset consisted of 84 entries, describing antibiotic resistance rates towards four classes of antimicrobial agents (Aminoglycosides, Aminopenicillins, Fluoroquinolones and third-generation Cephalosporins) across 21 countries. The effect sizes, associated standard errors and significance tests of the model parameters are presented in Table 1. Note the strong effect of antibiotic use in the hospital sector which is two to three orders of magnitude higher than the effect of the other two predictors. The effect of antibiotic usage in the veterinary medical sector on resistance rates is slightly below zero, therefore it is likely that the true correlation is equal to zero.

**Table 1:** Estimated effect sizes of the three fixed-effects predictors on antibiotic resistance rates in E. coli. Notice the significantly bigger effect of antibiotic use in the hospital sector compared to the two other effect sizes. The negative effect size of the third predictor is likely to be indicative of a true effect size of zero. Stars indicate statistical significance at the 0.05 (*), 0.01 (**) or 0.001 (***) level.

| Predictor | Estimate | SE | Z | p |
| --- | --- | --- | --- | --- |
| Hospital sector | $1.02 \times 10^{-1}$ | $4.843 \times 10^{-3}$ | 21.124 | $< 2 \times 10^{-16}$*** |
| Primary care sector | $9.72 \times 10^{-4}$ | $3.882 \times 10^{-4}$ | 2.505 | 0.012* |
| Veterinary medical sector | $-1.95 \times 10^{-3}$ | $4.669 \times 10^{-4}$ | -4.155 | $3.26 \times 10^{-5}$*** |

#### 3.2.2 K. pneumoniae

The final dataset consisted of 62 entries, describing antibiotic resistance rates towards three classes of antimicrobial agents (Aminoglycosides, Fluoroquinolones and third-generation Cephalosporins) across 21 countries. The effect sizes, associated standard errors and significance tests of the model parameters are presented in Table 2. Again, note the strong effect of antibiotic use in the hospital sector which is significantly higher than the effect of the other two predictors.

**Table 2:** Estimated effect sizes of the three fixed-effects predictors on antibiotic resistance rates in K. pneumoniae. Notice the significantly bigger effect of antibiotic use in the hospital sector compared to the two other effect sizes. Stars indicate statistical significance at the 0.05 (*), 0.01 (**) or 0.001 (***) level.

| Predictor | Estimate | SE | Z | p |
| --- | --- | --- | --- | --- |
| Hospital sector | $1.82 \times 10^{-1}$ | $3.641 \times 10^{-2}$ | 5.003 | $5.66 \times 10^{-7}$*** |
| Primary care sector | $1.85 \times 10^{-3}$ | $5.019 \times 10^{-3}$ | 0.368 | 0.713 *n.s.* |
| Veterinary medical sector | $1.33 \times 10^{-2}$ | $3.187 \times 10^{-3}$ | 4.175 | $2.97 \times 10^{-5}$*** |

### 3.3 Representative predictions

In order to illustrate the effects of the three predictors, we generated predictions and 0.65-confidence bands of resistance rates as function of antibiotic usage rates in the three examined sectors. The following figures present our results for resistance rates of the two bacterial species towards Fluoroquinolones for the country of Denmark. The same plots for another representative country, Belgium, are presented afterwards, as well as the plots for Denmark for another class of antibiotics, the third-generation Cephalosporins.

**Figure 1:**
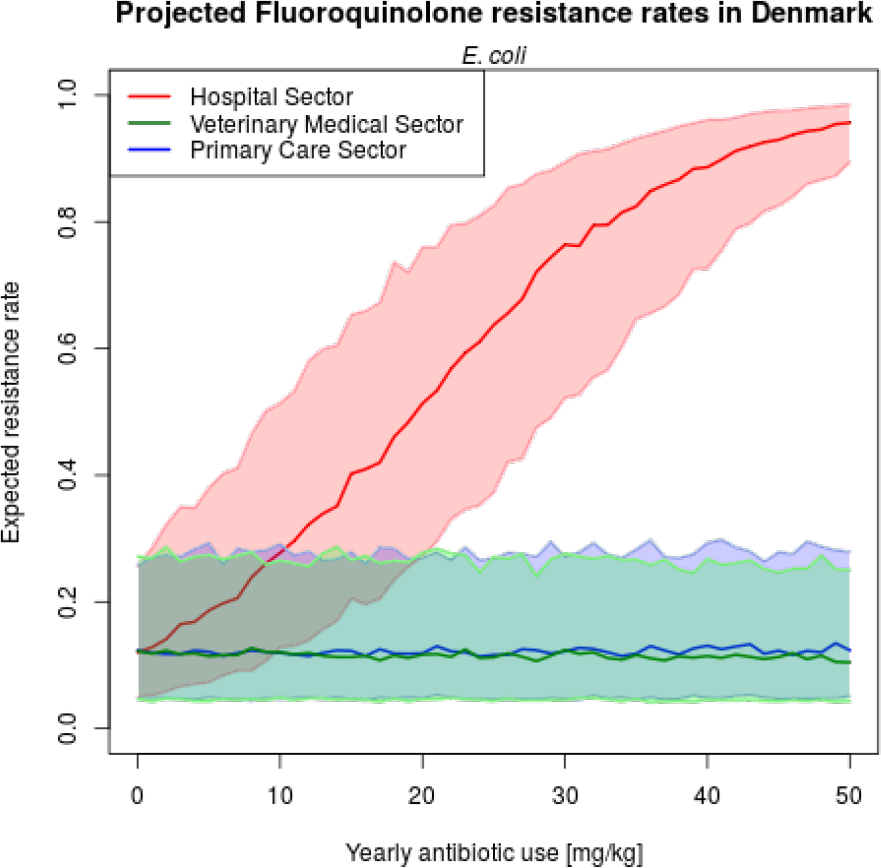
Predicted resistance rates of E. coli towards Fluroquinolones in Denmark as a function of yearly antibiotic use per sector.

**Figure 2:**
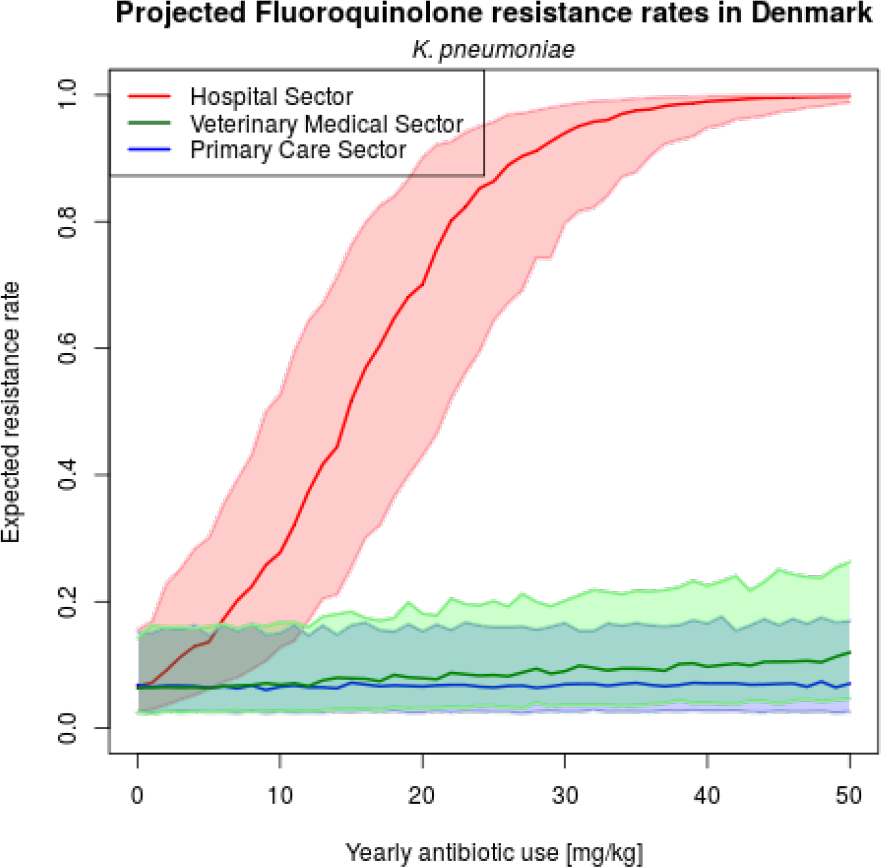
Predicted resistance rates of K. pneumoniae towards Fluroquinolones in Denmark as a function of yearly antibiotic use per sector.

**Figure 3:**
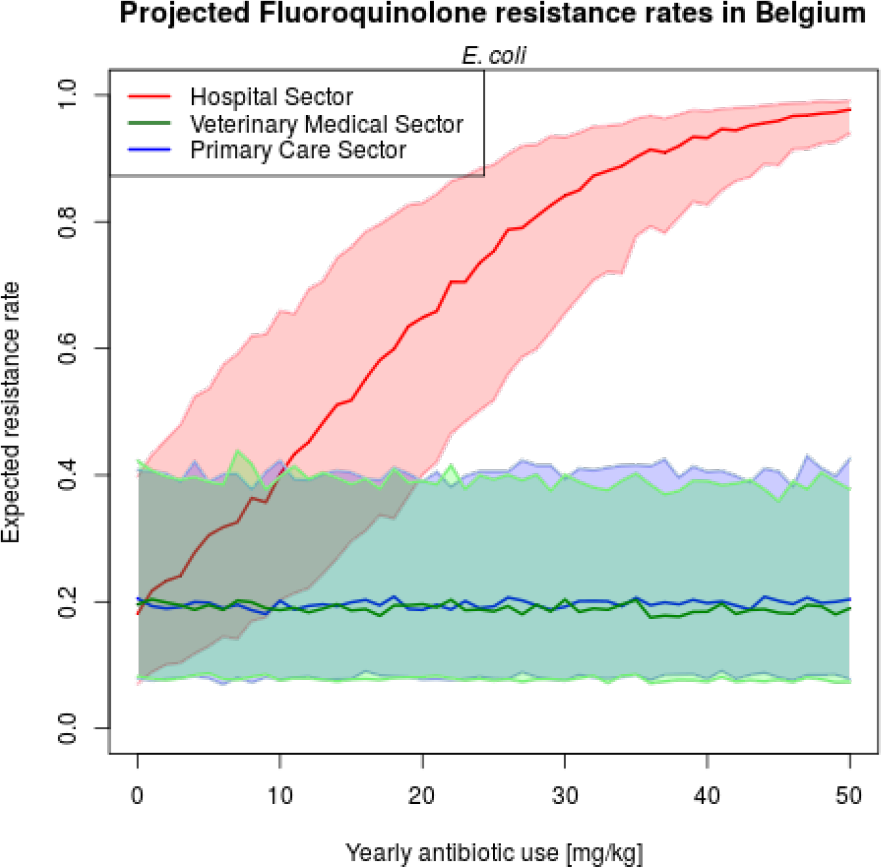
Predicted resistance rates of E. coli towards Fluroquinolones in Belgium as a function of yearly antibiotic use per sector.

**Figure 4:**
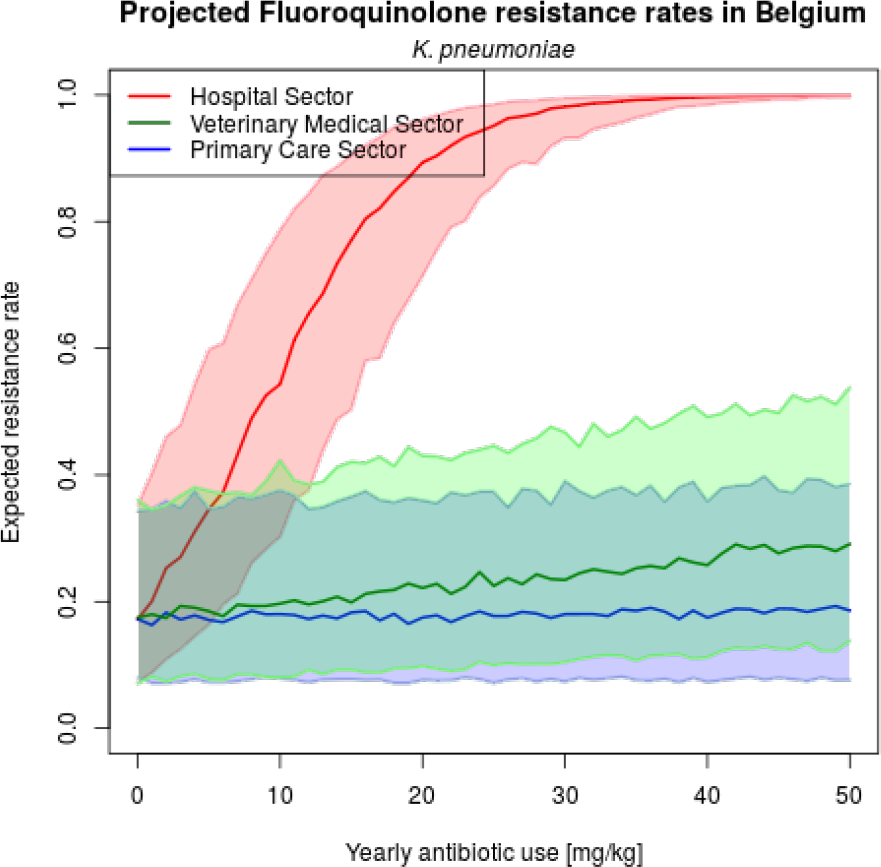
Predicted resistance rates of K. pneumoniae towards Fluroquinolones in Belgium as a function of yearly antibiotic use per sector.

**Figure 5:**
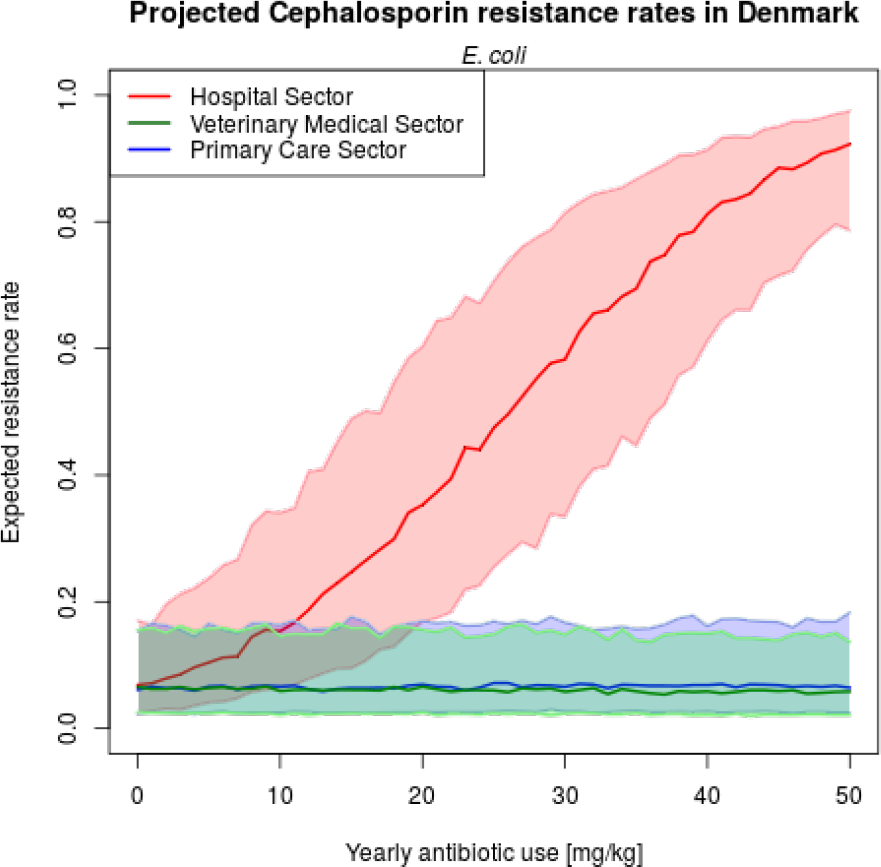
Predicted resistance rates of E. coli towards third-generation Cephalosporins in Denmark as a function of yearly antibiotic use per sector.

**Figure 6:**
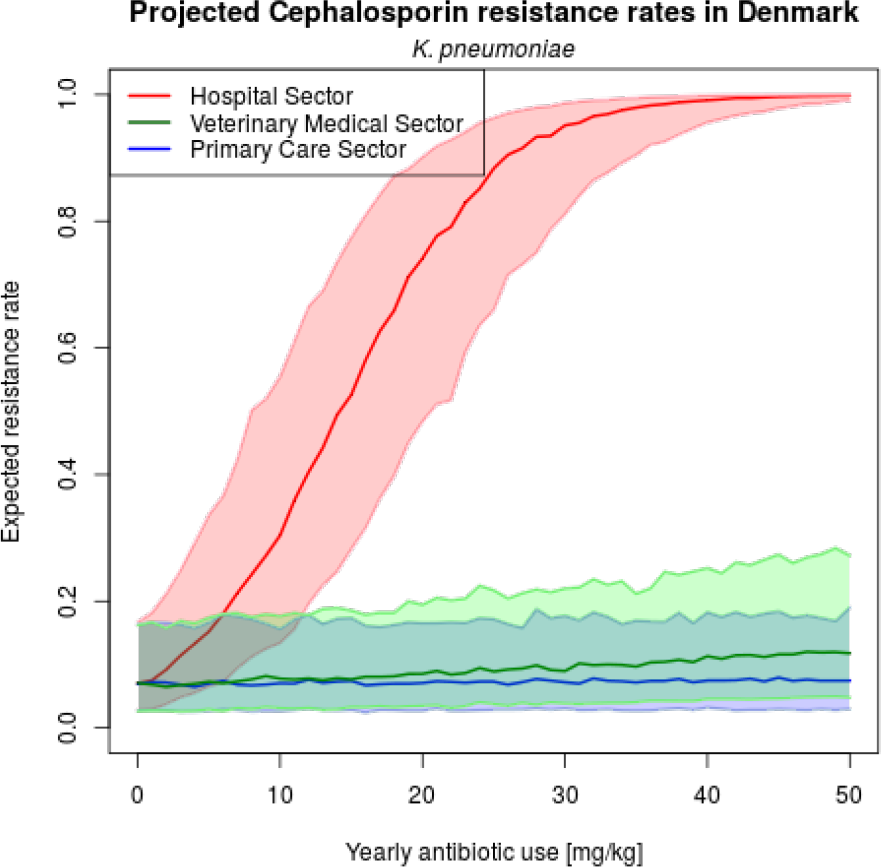
Predicted resistance rates of K. pneumoniae towards third-generation Cephalosporins in Denmark as a function of yearly antibiotic use per sector.

## 4 Discussion

In this work, we have demonstrated a high correlation between antibiotic use in the hospital sector and population level rates of infections with antibiotic-resistant bacteria. Moreover, we have demonstrated that this effect is considerably larger than the effect of antibiotic use in the primary care sector and the effect of veterinary antimicrobial consumption. Our results fall in line with the theory of antibiotic resistance emerging under the selective pressure of wide-spread antibiotic use in humans. In contrast, the presented analyses seem to challenge the idea of veterinary medical antibiotic usage as a main driver of infections with antibiotic-resistant bacteria. The second interesting observation is the significantly lower effect of antibiotic use in the primary care sector compared to the effect of hospital antibiotic use.

The development and acquisition of resistance to antibiotic substances of microbes within agricultural facilities has been repeatedly demonstrated (Witte, 1998; Teuber, 2001; Heuer et al., 2011). It is, therefore, an interesting question why these resistant microbes do not seem to be able to jump to humans more frequently. One reason might be the fact that they usually encounter healthy humans with an intact immune system. Moreover, it can be assumed that the probability of resistant microbes to escape the food industry is probably rather low when hygiene standards during breeding, transport, slaughter and consumption of meat are respected.

The difference between the effects of the primary care and the hospital sector is, in fact, even more important than the previous finding and requires further investigation. At the current point of time, one can only carefully speculate on the reasons for this difference. Possible reasons could include the fact that hospitals concentrate a high number of diseased people in a small amount of space, thereby making the transmission of possibly antibiotic-resistant bacterial strains more likely. Moreover, the use of immunosuppressive drugs to prevent the rejection of transplanted organs or tissues and for treating inflammatory diseases makes infection with such bacteria more likely as well. Furthermore, the behaviour of hospital staff, be it medical or service personnel, might exacerbate this effect. Poor hand sanitation and a disregard of other hygiene standards, such as hospital staff fixing open the door to rooms of patients with airborne diseases so they can see the patient at all times, have been reported in the literature and are of serious concern (Eames et al., 2009). Altogether, the generally high numbers of hospital-associated infections and subsequent fatalities (Klevens et al., 2007) hint at serious problems in this sector of health care. At any rate, the flagrant differences between the hospital sector on the hand and the primary medical care and veterinary sector on the other hand call for thorough surveillance and enforcement of hygienic measures on the hospital sector.

The results presented are subject to a number of limitations. First, for reasons of statistical power, only the analysis of data on resistance rates of two bacterial species was possible, however both analyses yielded very similar results. Second, due to reasons of comparability, only data from the European Union was considered, a zone of rather homogeneous, and thereby meaningfully comparable countries, which has led to the drawback of a rather small data corpus. Third and as a consequence of this, the fitted statistical models had to assume the same slopes across the four (or, in case of *K. pneumoniae*, three) antibiotic classes in order to reduce the number of required degrees of freedom. While we believe this to be a reasonable assumption, a definite confirmation of this is still pending. To be sure, we have also fitted more complex models allowing for differing random slopes across the different antibiotic classes which yielded qualitatively similar results, but were of limited use because the high number of estimated parameters compared to the rather small data basis has led to a high statistical uncertainty of the obtained estimates. For the same reason of limited available data, we were also not able to estimate the effect sizes of possible interactions between the main effects. Finally, the reanalysis of observational data comes with the typical disadvantage of a much more limited mechanistic understanding of the involved processes compared to experimental studies carried out under controlled conditions.

Despite these limitations, we still believe these results to be an important starting point for further investigation with high relevance to the field of public health. With regard to theoretical examinations, we propose similar statistical analyses operating on bigger sets of data in order to enhance statistical power and allow for a more precise estimation of effect sizes. Extending the analyses to a broader range of countries is imaginable, as well as analyses operating on datasets spanning broader time scales. Furthermore, we consider spatial analyses of case data of infections with antibiotic-resistant bacterial strains helpful which specifically examine the question whether such cases show significant clustering around agricultural facilities. This way, both the effect size and also the radius of influence of such facilities on resistance rates could be quantified.

With regard to microbiological studies, we specifically propose the examination of the exact mechanisms leading to the strong influence of hospital antibiotic usage on resistance rates on the population level. Moreover, we propose analysing possible ways of transmission of resistant bacterial strains from the inside of livestock facilities into their environment if such a transmission has to be expected.

## 5 Data availabilty

The compiled dataset and the R script used for the analysis are available from the authors upon request and will be publicly available soon.

## 6 Author contributions

MMF conceived the study together with MB, extracted and analysed the data and wrote the manuscript. MB provided critical feedback to both the methodology and the initial draft of the manuscript and checked all statistical analyses. Both authors have read and approve of the final version of this work.

## 8 Acknowledgements

We express our gratitude towards Prof. R. Mutzel for his constant encouragement, helpful discussions and critical reading of the manuscript.

## 9 Appendix A: Diagnostic plots for model assumptions

**Figure 7:**
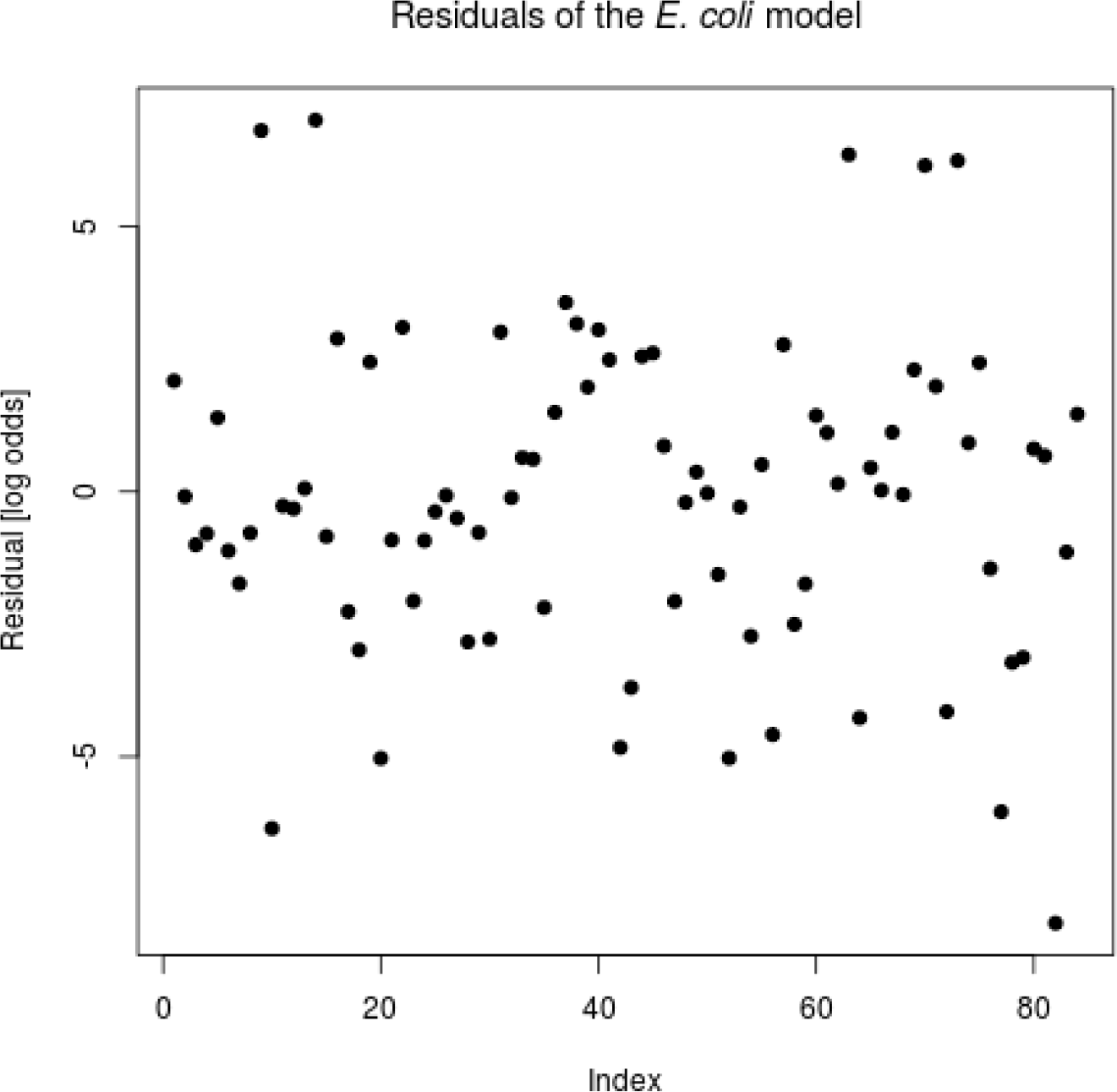
Diagnostic plot showing the residuals of the model fitted for the resistance rates of E. coli. Normality has been confirmed with a Shapiro-Wilk test (see results section).

**Figure 8:**
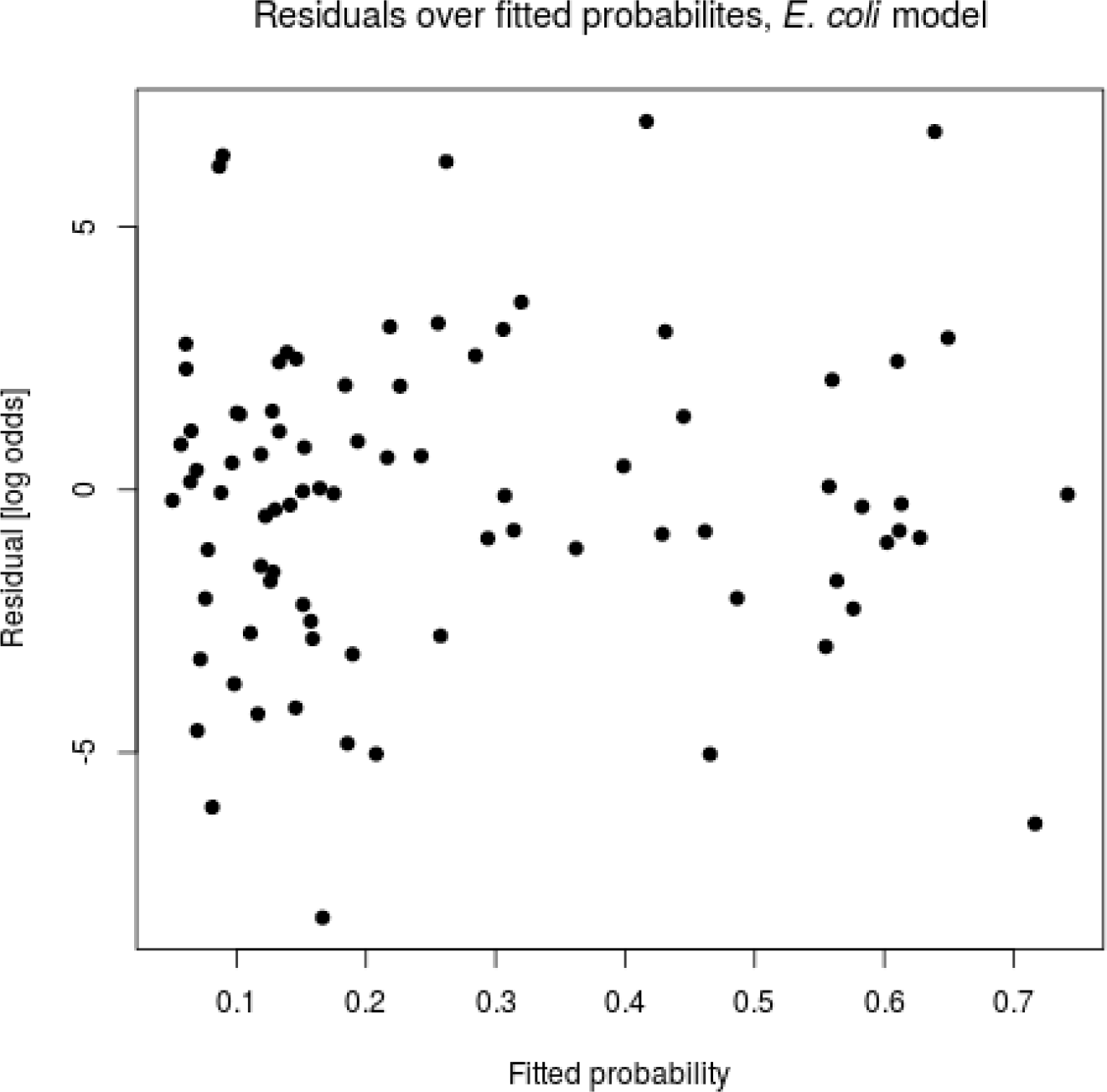
Diagnostic plot showing the residuals of the model fitted for the resistance rates of E. coli as a function of the fitted probabilities. Notice the approximately constant variance over the complete range of values indicating homoscedasticity.

**Figure 9:**
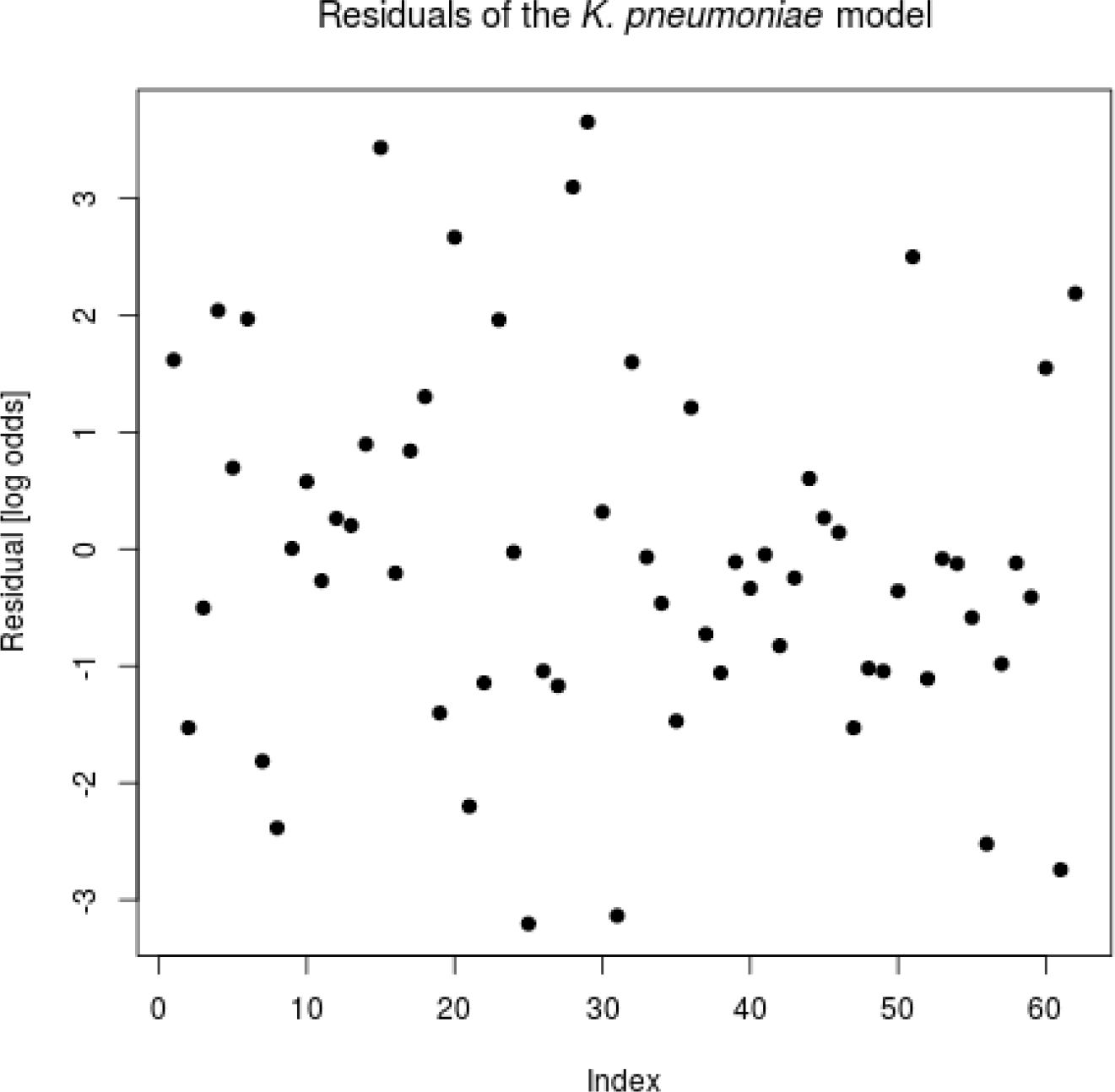
Diagnostic plot showing the residuals of the model fitted for the resistance rates of K. pneumoniae. Normality has been confirmed with a Shapiro-Wilk test (see results section).

**Figure 10:**
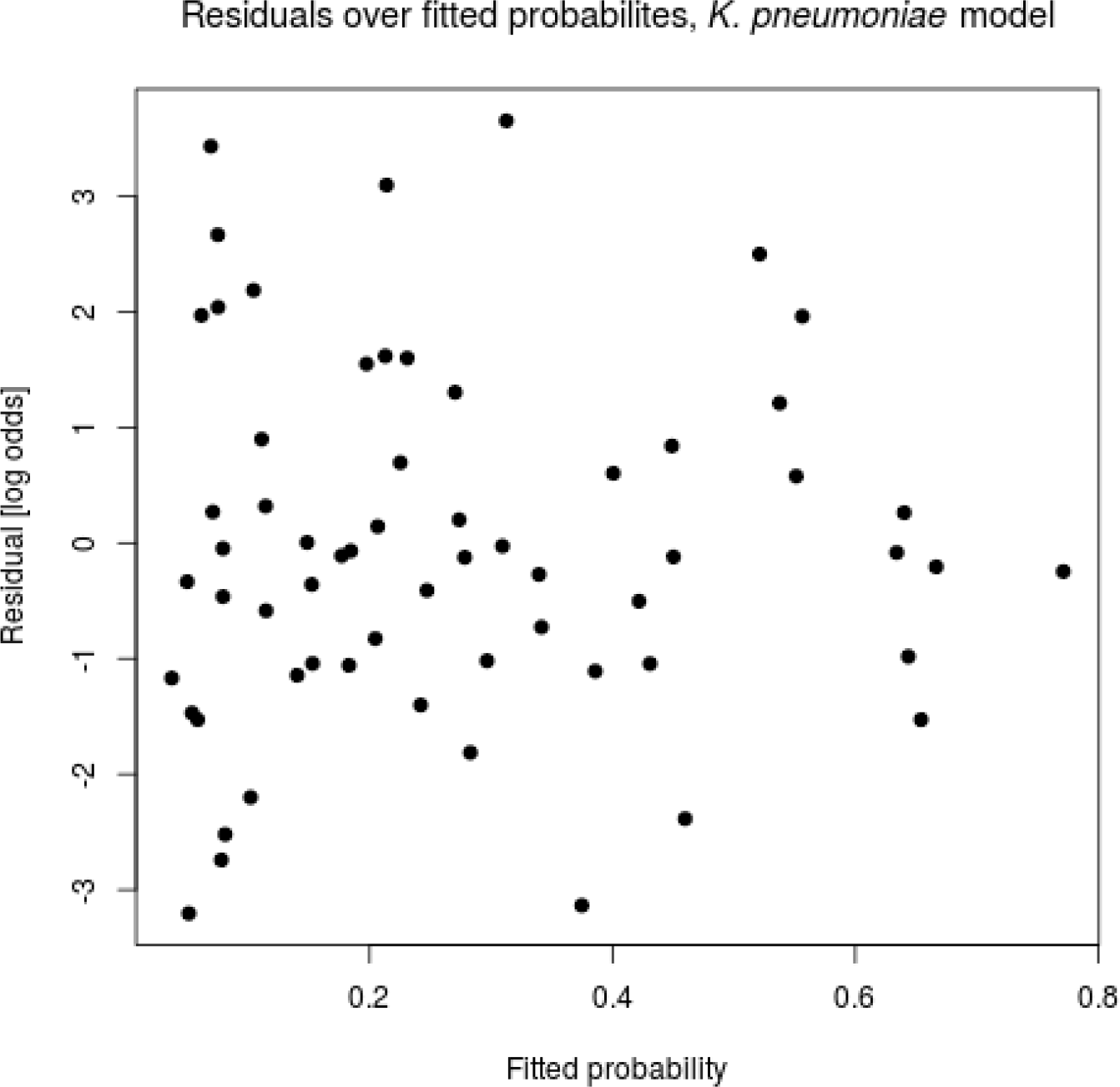
Diagnostic plot showing the residuals of the model fitted for the resistance rates of K. pneumoniae as a function of the fitted probabilities. Notice the approximately constant variance over the complete range of values indicating homoscedasticity.

## 10 Appendix B: Diagnostic plots for goodness of fit

**Figure 11:**
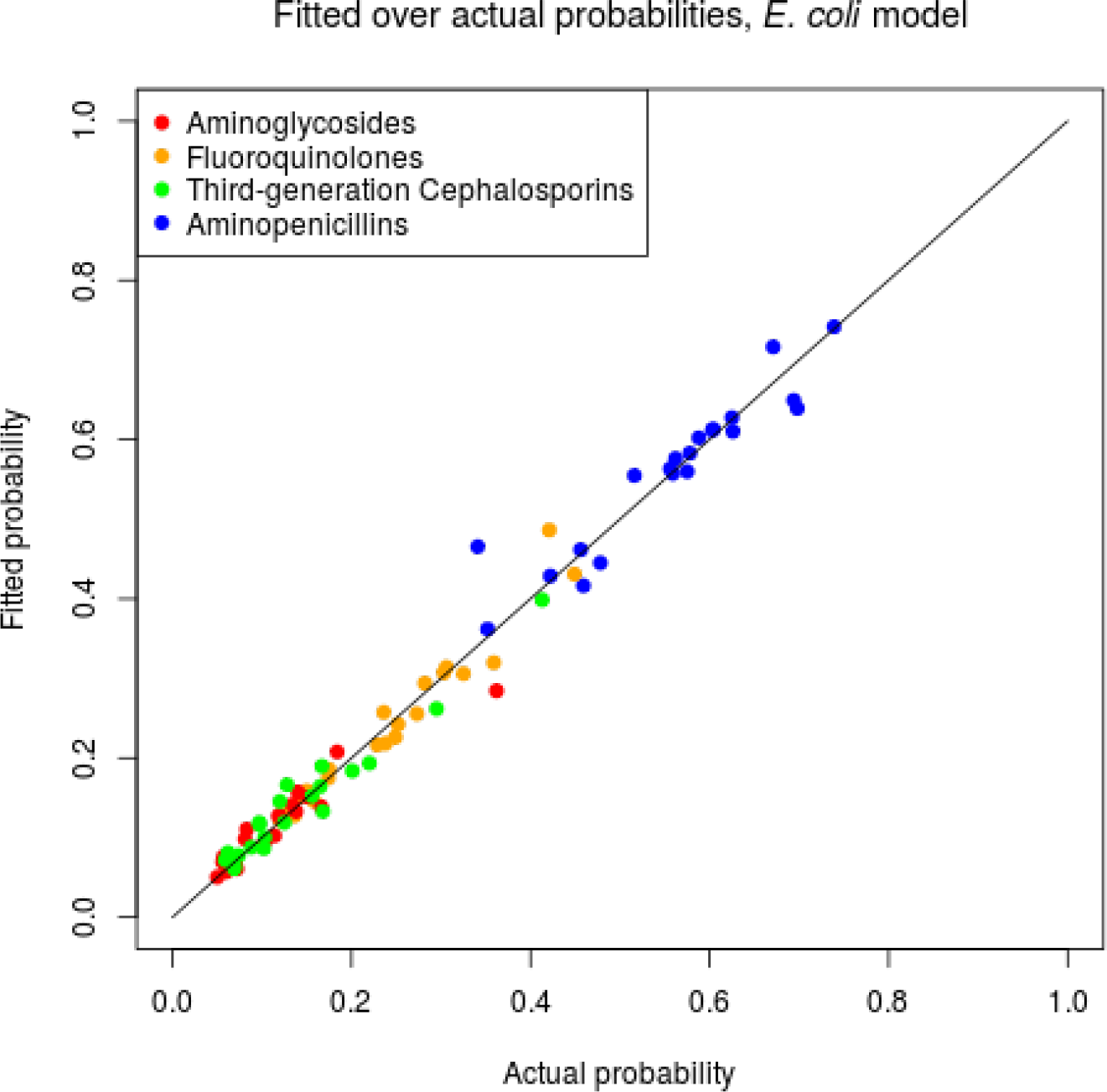
Diagnostic plot showing the fitted resistance rates of E. coli as a function of the actual ones. The continuous line represents the diagonal y = x. Notice the clear correlation indicating a high goodness of fit.

**Figure 12:**
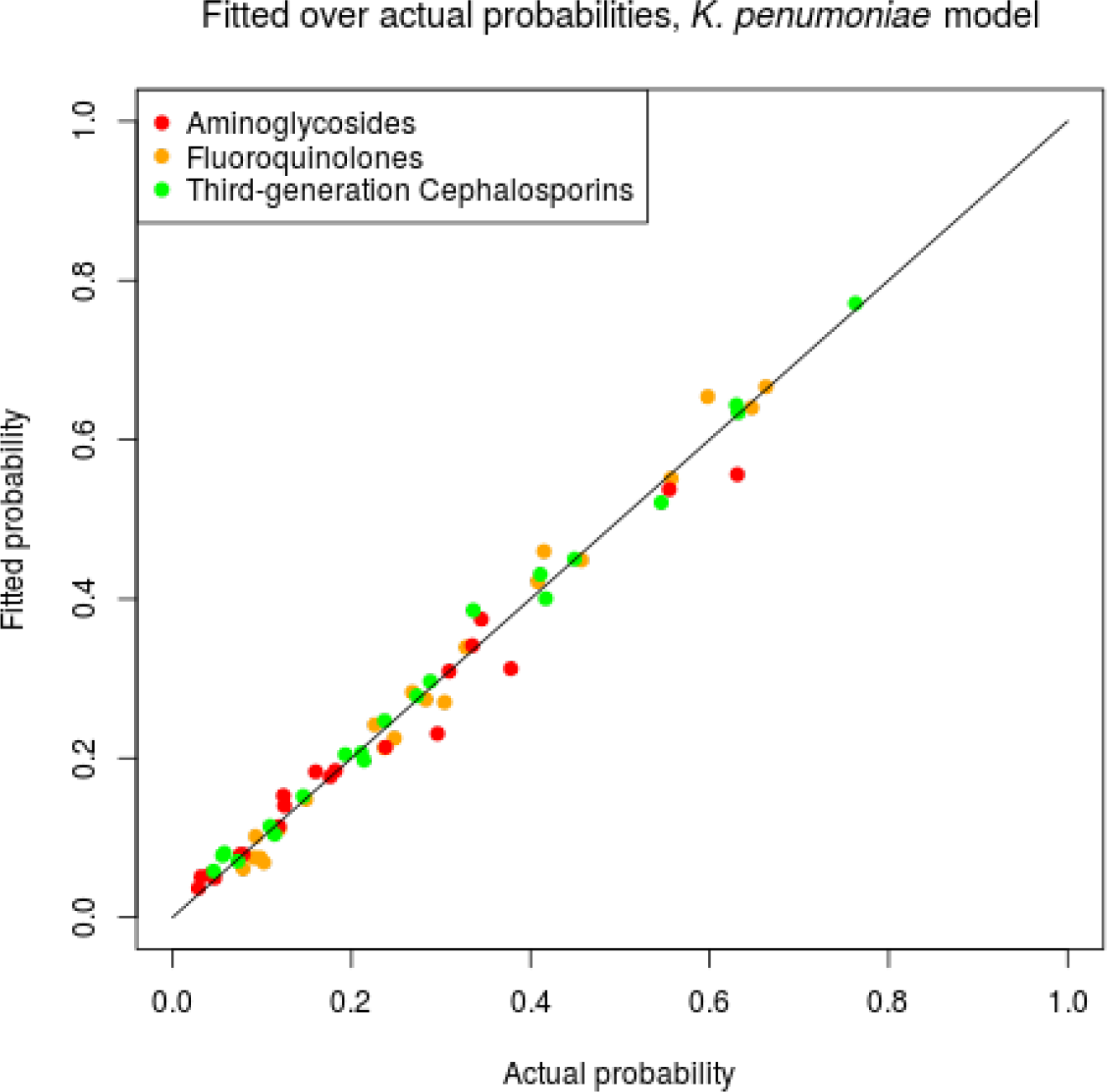
Diagnostic plot showing the fitted resistance rates of K. pneumoniae as a function of the actual ones. The continuous line represents the diagonal y = x. Notice the clear correlation indicating a high goodness of fit.

